# Mini-heterochromatin domains constrain the *cis*-regulatory impact of SVA transposons in human brain development and disease

**DOI:** 10.1101/2023.10.20.563233

**Authors:** Vivien Horváth, Raquel Garza, Marie E. Jönsson, Pia A. Johansson, Anita Adami, Georgia Christoforidou, Ofelia Karlsson, Laura Castilla Vallmanya, Patricia Gerdes, Ninoslav Pandiloski, Christopher H. Douse, Johan Jakobsson

## Abstract

SVA retrotransposons remain active in humans and contribute to individual genetic variation. Polymorphic SVA alleles harbor gene-regulatory potential and can cause genetic disease. However, how SVA insertions are controlled and functionally impact human disease is unknown. Here, we dissect the epigenetic regulation and influence of SVAs in cellular models of X-linked dystonia-parkinsonism (XDP), a neurodegenerative disorder caused by an SVA insertion at the *TAF1* locus. We demonstrate that the KRAB zinc finger protein ZNF91 establishes H3K9me3 and DNA methylation over SVAs, including polymorphic alleles, in human neural progenitor cells. The resulting mini-heterochromatin domains attenuate the *cis*-regulatory impact of SVAs. This is critical for XDP pathology; removal of local heterochromatin severely aggravates the XDP molecular phenotype, resulting in increased *TAF1* intron retention and reduced expression. Our results provide unique mechanistic insights into how human polymorphic transposon insertions are recognized, and their regulatory impact constrained by an innate epigenetic defense system.

## Introduction

More than 50% of the human genome is made up of transposable elements (TEs) ^1–3^. Three families of TEs are still active in humans: the autonomous long interspersed nuclear element 1 (LINE-1); the non-autonomous Alu short interspersed element (Alu); and the composite element SINE-R-VNTR-Alu (SVA) ^4–9^. The mobilization of these elements represents a significant source of genomic variation in the human population and is the underlying cause of some genetic diseases ^6,10–15^.

SVAs are a class of hominoid-specific TEs. They are non-autonomous, depending on the LINE1 machinery for retrotransposition, and consist of a fusion of two TE fragments separated by a variable number of tandem repeats (VNTR) ^16–19^. Based on their evolutionary age, SVAs are divided into different subfamilies (A-F), of which SVA-E and SVA-F are human-specific ^5^ and make up about half of the approximately 6000 fixed SVAs annotated in the human genome ^20^. In addition to the annotated SVAs, there are thousands of polymorphic SVA alleles in the human population. Current estimates suggest about one new germline SVA insertion in every 60 births ^5,10,21–24^. The individual genetic variation caused by polymorphic SVA insertions is thought to contribute to phenotypic variation in the human population and contribute to, or cause, disease ^11,12,25^. However, SVAs have been notoriously challenging to study due to their highly repetitive nature, and little is known about how polymorphic SVA insertions are regulated by the human genome or how they influence phenotypic traits and disease.

SVAs harbor strong gene regulatory sequences that can function both as transcriptional activators and repressors, influencing the expression of genes in the vicinity of their integration site ^25–32^. Notably, SVAs appear to be particularly potent as *cis*-regulatory elements in the human brain, where they have been linked to enhancer-like activities ^29,32^. In line with this, polymorphic SVAs have been linked to several genetic neurological disorders ^23,33–37^. The most well-characterized of these is X-linked dystonia-parkinsonism (XDP), a recessive adult-onset autosomal genetic neurodegenerative disorder ^38–40^. XDP is caused by a germline SVA retrotransposition event in intron 32 of *TAF1*, a gene that encodes TATA-box binding protein associated factor 1, an essential part of the transcriptional machinery ^39,41^. The SVA insertion interferes with the transcription and/or splicing of *TAF1* mRNA, resulting in reduced expression ^39,41^. The example of XDP illustrates the important role of polymorphic SVA insertions in human brain disorders. However, although the SVA insertion is the underlying genetic cause of XDP, the molecular mechanism behind how the SVA interferes with *TAF1* expression is unknown and there is still no mechanistic insight into why certain SVA insertions cause brain disorders. For example, there are hundreds of intronic SVA insertions in the human genome that do not cause disease. How is the human brain protected against the strong regulatory impact of SVAs in these cases and what makes the disease-causing SVA insertion in *TAF1* unique?

In this study we demonstrate that the DNA-binding KRAB zinc finger protein (KZFP) ZNF91 plays a key role in protecting the human genome against the *cis*-regulatory impact of SVAs by establishing a dual layer of repressive epigenetic modifications over SVAs in neural cells, including new polymorphic alleles such as the disease-causing XDP-SVA. The resulting mini-heterochromatin domains are characterized by the presence of both DNA methylation and H3K9me3. Notably, the presence of ZNF91-mediated heterochromatin on the polymorphic XDP-SVA is highly relevant for XDP pathology, as the removal of this heterochromatin domain aggravates the molecular XDP phenotype, resulting in increased intron-retention and reduced *TAF1* expression. In summary, our results provide unique mechanistic insights into how human polymorphic TE insertions are recognized, and how their potential regulatory impact in neural cells is minimized by an innate epigenetic defense system based on a KZFP.

## Results

### Establishing XDP-NPCs to study the epigenetic regulation of SVAs

To investigate the molecular mechanisms controlling SVAs in human neural cells, including the polymorphic XDP-SVA, we established a neural progenitor cell (NPC) model system using induced pluripotent stem cell (iPSC) lines derived from three XDP patients and three control individuals (Figure 1A, Table 1). The XDP-SVA carriers presented initially with dystonia at a mean age at onset of 42.6 years (±13.6), similar to what has previously been reported (42.3±8.3 years) (Table 1) ^39,42^. The controls used were unaffected sons of two of the XDP-SVA carriers (Table 1). The six iPSC lines were converted into stable NPC lines ^43^ (XDP- and Ctrl-NPCs) that could be extensively expanded or differentiated into different neural cell types. The XDP- and Ctrl-NPCs exhibited NPC morphology and expressed NPC markers such as SOX2 and NESTIN, monitored with immunocytochemistry (Figure 1B, Figure S1A). The expression of NPC markers, as well as the lack of expression of pluripotency markers, was also confirmed by RNA-seq (Figure 1C).

**Figure 1.**
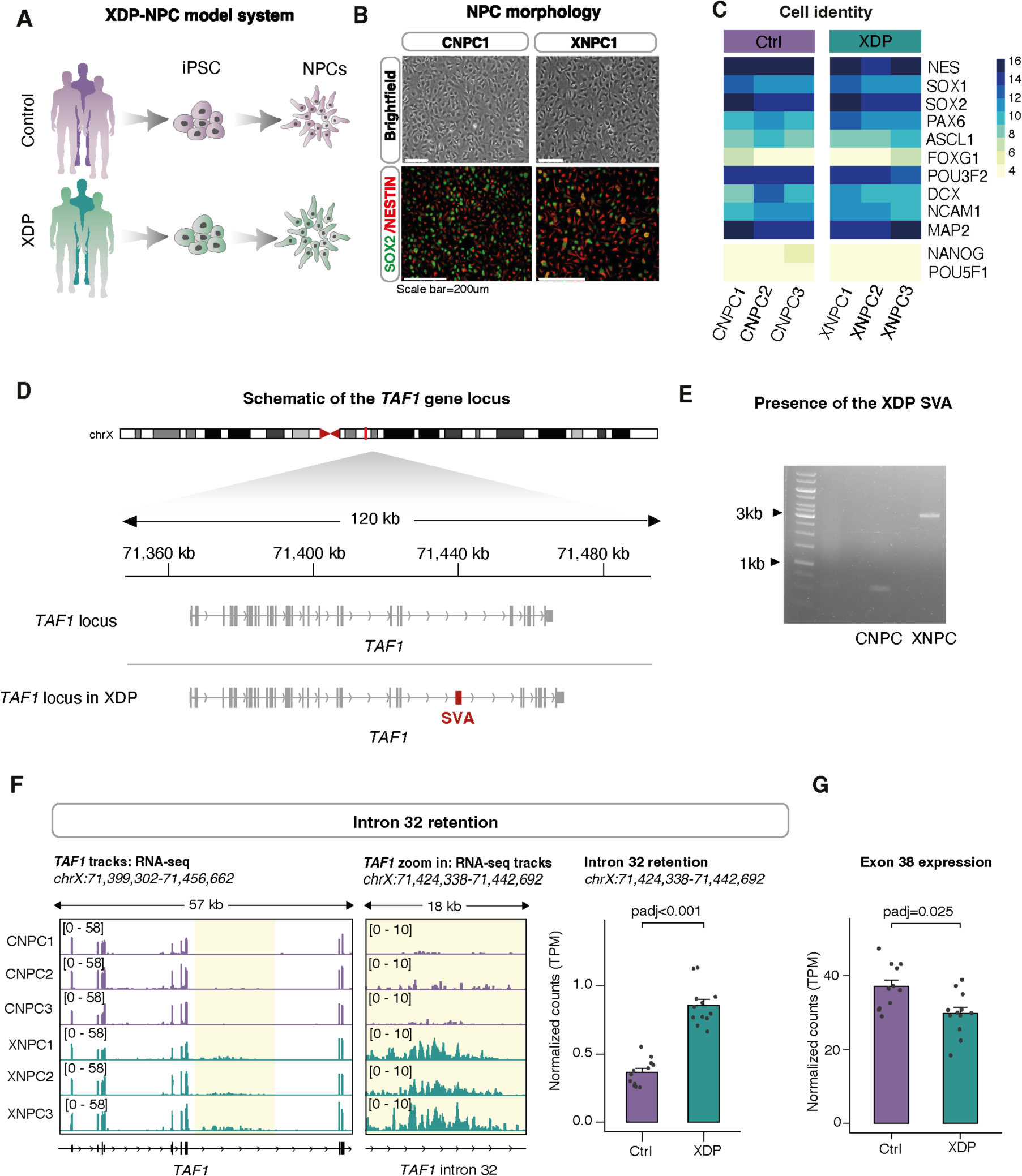
Characterization of the XDP-NPC model system. (A) Schematic of the generation of XDP-NPCs. (B) Brightfield images of Ctrl-(CNPC1) and XDP-NPCs (XNPC1) (top). Immunocytochemistry (bottom) of Sox2 (green) and nestin (red) in Ctrl- and XDP-NPCs. (C) Heatmap of NPC marker-gene expression in Ctrl-NPCs (n=3) and XDP-NPCs (n=3) measured using RNA-seq. (D) Schematic of the *TAF1* gene locus. The polymorphic XDP-SVA is depicted in red. (E) PCR analysis of genomic DNA identifying the XDP-SVA. (F) Genome browser tracks showing gene expression of the *TAF1* gene (left) and a magnification of intron 32 of *TAF1*, highlighting the characteristic intron retention in XDP-NPCs. Quantification of *TAF1* intron 32 retention (right) in Ctrl-(n=12) and XDP-NPCs (n=12) (padj, DESeq2). (G) Quantification of *TAF1* exon 38 expression in Ctrl-(n=12) and XDP-NPCs (n=12) (padj, DESeq2).

**Table 1.**

| Status | Subject | Sample ID | Age at onset (years) | Age at collection (years) | Relationship |
| --- | --- | --- | --- | --- | --- |
| XDP | 33363.C | XNPC 1 | 38 | 44 | Father of CNPC 1 |
|  | 33109.2B | XNPC 2 | 58 | 72 | Father of CNPC 2,3 |
|  | 32517.B | XNPC 3 | 32 | 35 | Not related |
| Control | 33362.C | CNPC 1 | - | 18 | Son of XNPC 1 |
|  | 33114.C | CNPC 2 | - | 34 | Son of XNPC 2 |
|  | 33113.2I | CNPC 3 | - | 42 | Son of XNPC 2 |

Subjects as described in Ito et al. 2016 <sup>94</sup>

The presence of the XDP-SVA insertion, which is ∼2.6 kbp long and located in intron 32 of the *TAF1* gene, was confirmed using PCR (Figure 1D, E). RNA-seq analysis confirmed that XDP-NPCs displayed a characteristic retention of intron 32 of *TAF1* (p-value<0.001, DESeq2) and lower expression of downstream *TAF1* exons, such as exon 38, when compared to Ctrl-NPCs (p=0.025, DESeq2) (Figure 1F, G). These observations are similar to those previously described for XDP-iPSC and NPC lines^39^.

### SVAs are covered by H3K9me3 in NPCs

TEs, including SVAs, are associated with heterochromatin in somatic tissues that correlate with their transcriptional silencing, and which may impact their regulatory potential ^44^. We chose to characterize the repressive histone mark H3K9me3, which is linked to heterochromatin, in fetal human forebrain tissue, two XDP-NPCs, and two Ctrl-NPCs using CUT&RUN analysis (Figure 2A). The computational analysis of histone marks on SVAs using CUT&RUN data is challenging due to their repetitive nature. This results in a large proportion of ambiguous reads. To avoid false conclusions due to multi-mapping artefacts, we used a strict unique mapping approach to investigate individual SVA elements (Figure 2A). With this bioinformatic approach, it is only possible to investigate the epigenetic status of the flanking regions of the SVAs where the unique genomic context allows us to discriminate reads without ambiguity, and the epigenetic modification can be traced to unique loci in the human genome. The boundaries of nearly all SVAs (>1 kbp in length) of the different subfamilies (A-F), both in the developing human forebrain and in NPCs, were enriched with H3K9me3 (Figure 2B, C, Figure S2A). However, the genomic context did not enable us to analyze the XDP-SVA with this approach. To resolve this issue, we developed a qPCR-based technique in combination with CUT&RUN (Figure 2A, see Materials and methods). This showed a clear enrichment of H3K9me3 at the boundary of the XDP-SVA in XDP-NPCs (Figure 2D, Figure S2C).

**Figure 2.**
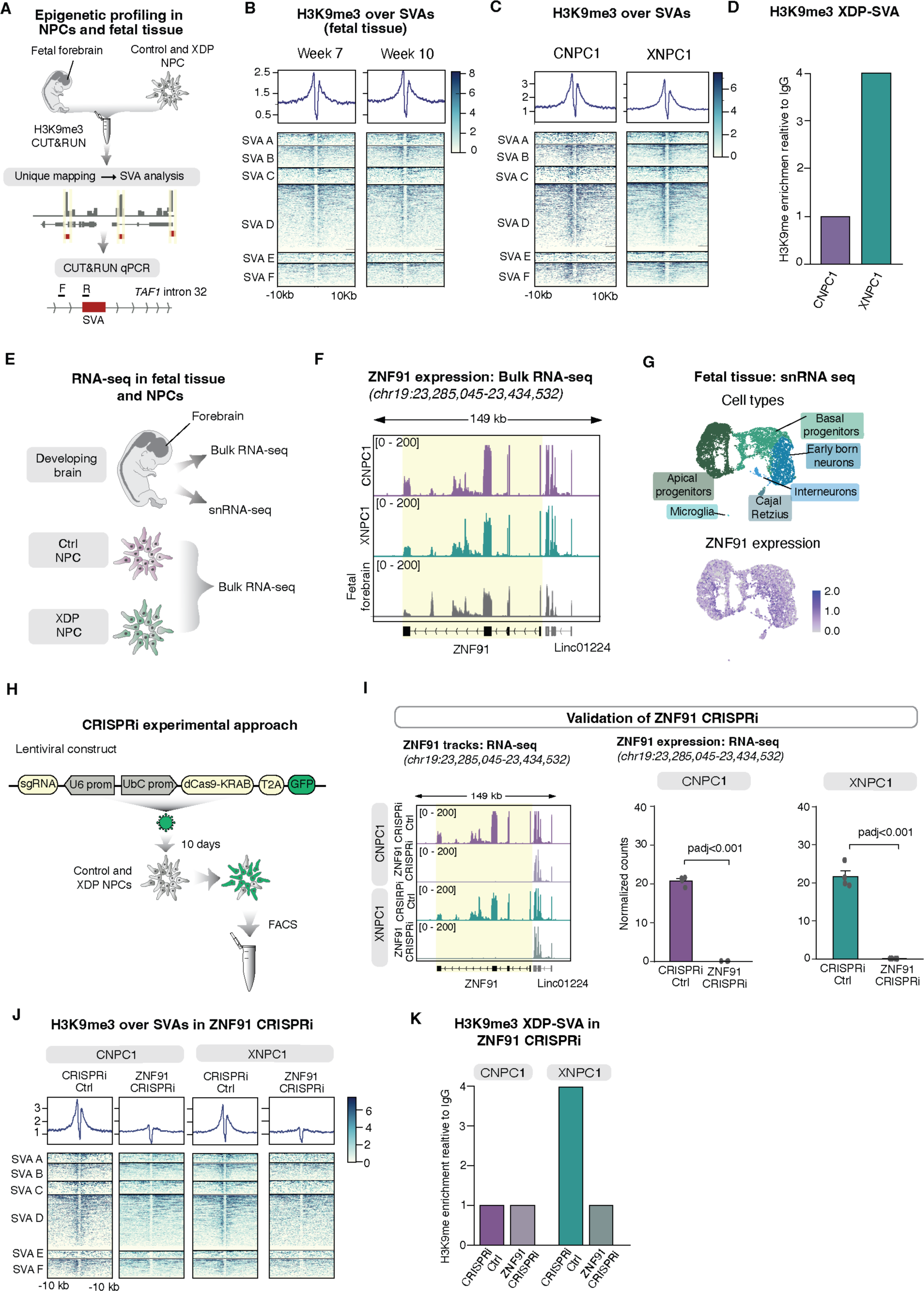
ZNF91 is required for H3K9me3 maintenance at SVAs in NPCs. (A) Schematic of CUT&RUN approaches to profiling H3K9me3 at SVAs in NPCs and human fetal forebrain tissue. (B) Heatmap showing enrichment of H3K9me3 over SVAs in human fetal forebrain tissue. (C) Heatmap showing H3K9me3 enrichment in NPCs. Displayed are the genomic regions spanning ±10 kbp up and downstream from the element. (D) Barplots showing the enrichment of H3K9me3 over the XDP-SVA and the lack of enrichment in control samples. (E) Schematic of RNA-seq and snRNA-seq experiments in NPCs and human fetal forebrain tissue. (F) RNA-seq tracks of *ZNF91* expression in NPCs and fetal forebrain. (G) UMAP showing characterized cell types (top). UMAP representing ZNF91 expression (bottom) in different cell types in the fetal brain. (H) Schematic of the CRISPRi approach including the lentiviral construct and experimental design. (I) RNA-seq tracks (left) and quantification (right) of *ZNF91* expression in CRISPRi-Ctrl and ZNF91-CRISPRi in Ctrl-NPCs (n=4) and XDP-NPCs (n=4) (padj, DESeq2). (J) Heatmap showing H3K9me3 over SVAs in CRISPRi-Ctrl and ZNF91-CRISPRi in Ctrl-NPCs and XDP-NPCs. (K) Barplots showing the effect of ZNF91-CRISPRi on H3K9me3 over the XDP-SVA in XDP-NPC.

### H3K9me3 deposition at SVAs is dependent on the KZFP ZNF91

To protect genomic integrity against TE insertions, organisms have evolved cellular defense mechanisms ^45,46^. KZFP genes have amplified and diversified in mammalian species in response to transposon colonization ^47–52^; recent profiling efforts have identified several KZFPs that bind to SVAs, including ZNF91 and ZNF611 ^29,50,53^. We noted that *ZNF91* is highly expressed in human fetal forebrain tissue and XDP- and Ctrl-NPC cultures, as monitored by bulk and snRNA-seq (Figure 2E-G) ^54^. Thus, we hypothesized that ZNF91 could be a KZFP that binds SVAs and recruits the epigenetic machinery that deposits H3K9me3 at these sites in NPCs.

To investigate a role for ZNF91 in SVA repression in NPCs, we designed a lentiviral CRISPR inhibition (CRISPRi) strategy to silence *ZNF91* expression. We targeted two guide RNAs (gRNAs) to a genomic region located next to the *ZNF91* transcription start site (TSS) and co-expressed gRNAs with a KRAB transcriptional repressor domain fused to catalytically-dead Cas9 (dCas9) (Figure 2H). As a control, we used a gRNA targeting lacZ, representing a sequence not found in the human genome. The transduction of XDP- and Ctrl-NPCs resulted in efficient silencing of *ZNF91*-expression, monitored with RNA-seq (Figure 2I, Figure S2D). CUT&RUN analysis of ZNF91-CRISPRi NPCs (XDP- and Ctrl-NPCs) revealed almost complete loss of H3K9me3 around SVAs (Figure 2J). This finding was reproduced in one additional XDP-NPC line and one additional Ctrl-NPC line (Figure S2E). CUT&RUN qPCR confirmed that the XDP-SVA also lost H3K9me3 in a ZNF91-dependent manner in XDP-NPCs (Figure 2K). Using a similar CRISPRi strategy we also confirmed that the H3K9me3 at SVAs, including the XDP-SVA, also depend on TRIM28, an epigenetic corepressor protein that is essential for the repressive action of KZFPs, in Ctrl- and XDP-NPCs (Figure S2F-H) ^47,55^. Together, these results demonstrate that a ZNF91/TRIM28-dependent mechanism establishes local H3K9me3 heterochromatin over SVAs in human NPCs, including the polymorphic disease-causing XDP-SVA.

### SVAs are covered by DNA methylation in human NPCs

In addition to H3K9me3, TE silencing in somatic tissues has also been extensively linked to DNA CpG-methylation ^46,56–59^. To investigate the presence of DNA methylation on SVAs in NPCs, we performed genome-wide methylation profiling using Oxford Nanopore Technologies (ONT) long-read sequencing (Figure 3A) ^60,61^ on one XDP-NPC line (XNPC1) and one Ctrl-NPC line (CNPC1). The long-read DNA methylation analysis revealed that the SVA elements of different subfamilies (A-F), which are CG-rich sequences, were all heavily methylated in human NPCs (Figure 3B). In addition, the polymorphic XDP-SVA was fully covered by DNA methylation (Figure 3C). Furthermore, we performed Cas9-targeted ONT sequencing over the XDP-SVA on the ZNF91-CRISPRi NPCs and CRISPRi-Ctrl XDP-NPCs (Figure 3D). These results demonstrated that the XDP-SVA was fully methylated in both the ZNF91-CRISPRi and CRISPRi-Ctrl XDP-NPCs (Figure 3E). Thus, SVAs in human NPCs, including the XDP-SVA, are covered by both DNA methylation and H3K9me3. Our results also indicate that the presence of DNA methylation at SVAs is not dependent on ZNF91-binding or H3K9me3 in this cell type.

**Figure 3.**
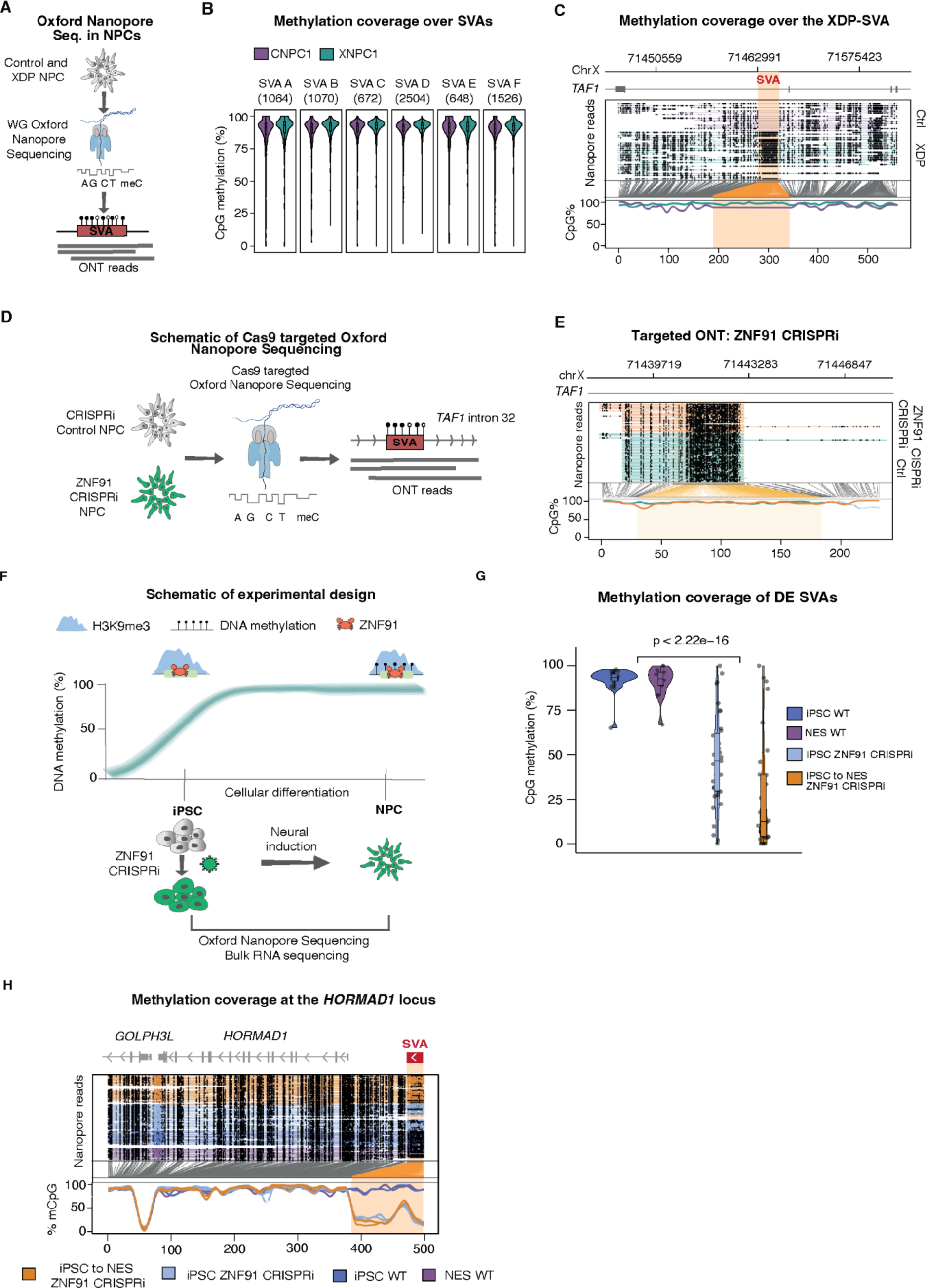
SVAs are covered by DNA methylation in NPCs. (A) Schematic of ONT sequencing experiment to monitor DNA methylation over SVAs. (B) Methylation coverage over SVAs in Ctrl- and XDP-NPCs. The different SVA families (A-F) are shown. (C) Methylation coverage over the XDP-SVA in Ctrl- and XDP-NPCs. (D) Schematic of Cas9 targeted ONT sequencing. (E) Targeted ONT sequencing in CRISPRi-Ctrl and ZNF91-CRISPRi NPCs. The *TAF1* XDP-SVA locus is shown. (F) Schematic of DNA methylation patterns during development in iPSCs and NPCs. ZNF91-CRISPRi in iPSCs and conversion to NPCs is also shown. (G) Violin plot showing DNA methylation over the first quarter (from their transcription start site) of the differentially-expressed SVAs (p-value, Student’s t-test). (H) DNA methylation pattern over an SVA element near the *HORMAD*1 gene.

### ZNF91 establishes DNA methylation on SVAs during early human development

Since the presence of DNA methylation on SVAs in NPCs did not depend on ZNF91, we wondered how and when DNA methylation on SVAs is established. DNA methylation is reprogrammed during the first few days of early development where global DNA methylation patterns, including that of many TEs, are erased and reinstated ^62–66^. During this process, TEs are initially silenced by dynamic epigenetic mechanisms which are then gradually replaced by other ^67^, more stable epigenetic mechanisms in somatic cell types such as NPCs ^55,68–71^. We and others have implicated TRIM28-KZFP complexes in this process ^55,69,71–75^. Thus, we hypothesized that ZNF91/TRIM28 may be involved in dynamically establishing DNA methylation of SVAs during earlier phases of human embryonic development (Figure 3F).

To test this hypothesis, we used iPSCs that resemble the epiblast stage of early human development^76^, where DNA methylation patterns are more dynamically regulated (Figure 3F). In contrast, NPCs are somatic cells with a stably methylated genome ^59,66,77^. We found clear evidence of dynamic ZNF91-mediated DNA methylation patterning of SVAs when we generated ZNF91-CRISPRi iPSCs. By performing genome-wide ONT analysis we found numerous SVAs (n=39) where DNA methylation was lost upon inhibition of ZNF91 (Figure 3G). In contrast, the same SVAs were covered by DNA methylation in control iPSCs and NPCs (Figure 3G). Notably, when we differentiated the ZNF91-CRISPRi iPSCs into NPCs we found that these SVAs remained hypomethylated (Figure 3G). Thus, without *ZNF91* expression DNA methylation could not be established on these SVAs upon differentiation. For example, an SVA-F element located upstream of the *HORMAD1* gene was fully methylated in control NPCs and iPSCs. The DNA methylation over this SVA was completely lost in ZNF91-CRISPRi iPSCs, and remained absent when the ZNF91-CRISPRi iPSCs were differentiated into NPCs (Figure 3H). These results demonstrate that the DNA methylation patterns over some SVAs are dynamic in iPSCs and depend on ZNF91. In addition, ZNF91 is essential for establishing the stable layer of DNA methylation found over these SVAs in NPCs. Thus, cellular context is important for the downstream consequence of ZNF91 binding to SVAs. In early development, ZNF91 mediates the establishment of both H3K9me3 and DNA methylation, while in somatic cells only H3K9me3 depends on ZNF91; DNA methylation is propagated through other mechanisms.

### DNA methylation and H3K9me3 co-operate to silence SVA expression in NPCs

To investigate the role of H3K9me3 and DNA methylation in the transcriptional silencing of SVAs in NPCs we combined loss-of-function experiments with RNA-seq analysis. To remove H3K9me3 we used the ZNF91-CRISPRi NPCs. To remove DNA methylation we deleted DNA methyltransferase 1 (*DNMT1*), which is the enzyme that maintains DNA methylation during cell division ^78^. We used a previously-described CRISPR-cut approach, resulting in a global loss of DNA methylation including over SVAs ^59,79^ as well as a CRISPRi combination strategy targeting the expression of both *DNMT1* and *ZNF91* in XDP- and Ctrl-NPCs (Figure 4A). Both approaches resulted in a global loss of DNA methylation, as monitored with 5mC immunocytochemistry (Figure 4B, Figure S3A), and a loss of DNA methylation over the XDP-SVA as demonstrated by targeted ONT long-read sequencing methylation analysis (Figure 4C). CUT&RUN analysis on DNMT1-KO NPCs revealed that loss of DNA methylation did not affect the presence of H3K9me3 at SVAs (Figure S3B).

**Figure 4.**
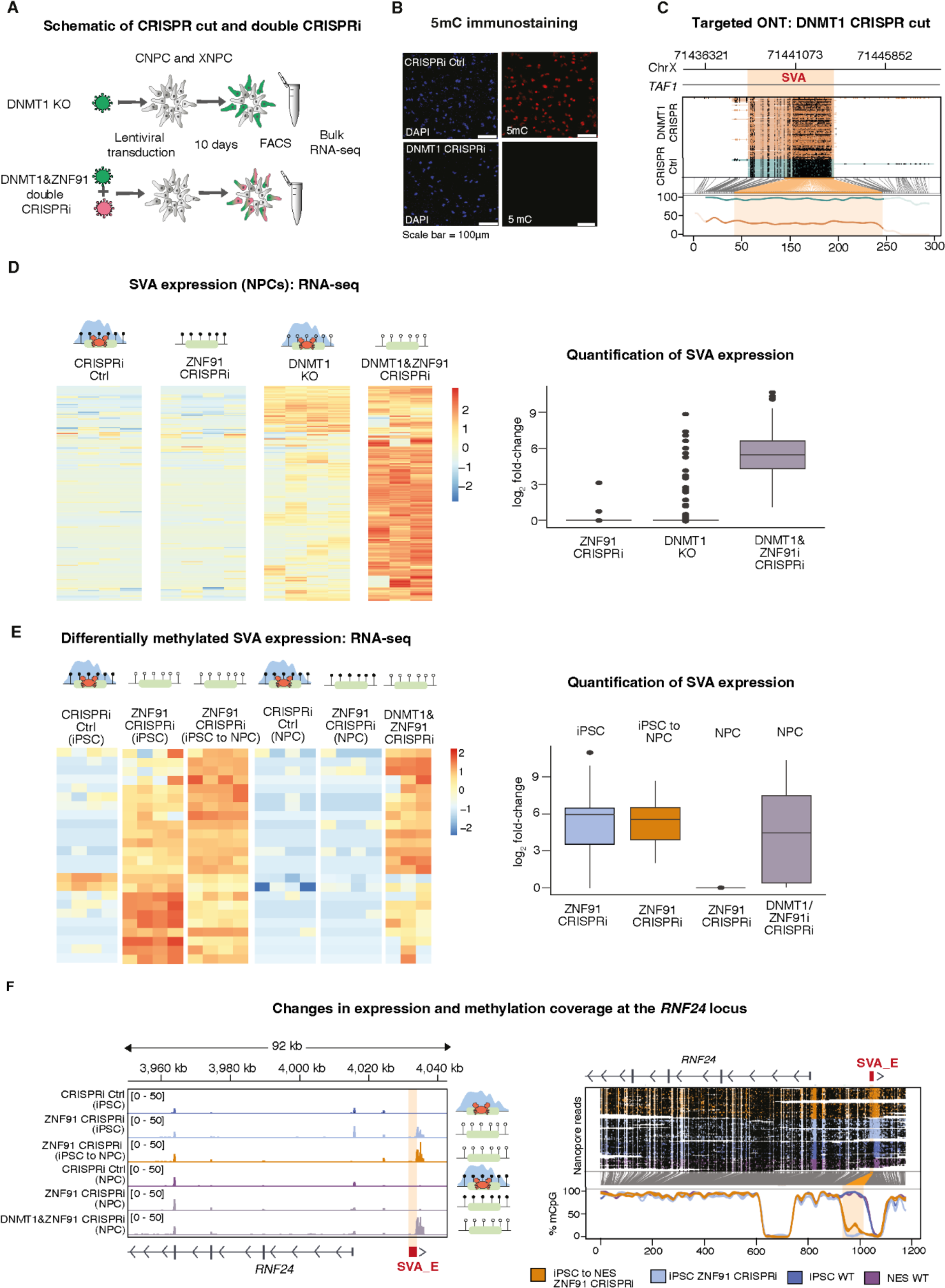
DNA methylation and H3K9me3 co-operate to silence SVAs in NPCs. (A) Schematic of CRISPR cut and double CRISPRi experiment in NPCs. (B) 5mC immunostaining showing the global loss of DNA methylation upon DNMT1-CRISPRi. (C) DNA methylation coverage over the XDP-SVA in CRISPR-Ctrl and DNMT1 CRISPR-cut conditions. (D) Heatmap (left) showing upregulated SVAs in ZNF91&DNMT1-CRISPRi. The same SVAs are also shown in ZNF91-CRISPRi and DNMT1-KO experiments. Boxplot (right) showing SVA expression in ZNF91-CRISPRi, DNMT1-KO, and ZNF91&DNMT1-CRISPRi. I Heatmap (left) showing the expression level of differentially-methylated SVAs. Boxplot (right) showing the expression of differentially-expressed SVAs. (F) Genome browser tracks and DNA methylation pattern over an SVA element near to the *RNF24* gene.

We used an in-house 2×150bp, polyA-enriched stranded library preparation for bulk RNA-seq using a reduced fragmentation step to optimize the read length for SVA analysis. Such reads can be uniquely assigned to many SVA loci. We obtained ∼40 million reads per sample. To quantify SVA expression we discarded all ambiguously mapping reads and only quantified those that map uniquely to a single location (unique mapping) ^80^. We found that ZNF91-CRISPRi in NPCs, which removes H3K9me3 at SVAs, did not result in activation of SVA expression (Figure 4D). When DNMT1 was deleted in NPCs, which removes DNA methylation, we also found only a small number of SVAs transcriptionally upregulated (Figure 4D). However, in the ZNF91&DNMT1-CRISPRi NPCs, where both H3K9me3 and DNA methylation over SVAs is lost, we found a massive transcriptional activation of hundreds of SVAs (Figure 4D).

We also analyzed the expression of SVAs in the iPSC-NPC conversion experiments (Figure 3F). RNA-seq revealed that the SVAs that lost DNA methylation after ZNF91 deletion in iPSCs (ZNF91-CRISPRi iPSCs, Figure 3G) were also transcriptionally upregulated (Figure 4E). This contrasted with ZNF91-CRISPRi NPCs, where the same SVA elements were not upregulated upon inhibition of ZNF91 (Figure 4E). When we analyzed the ZNF91-CRISPRi iPSCs that were differentiated to NPCs we found that the SVAs were expressed in these NPCs (Figure 4E, F). These SVAs were also found to be upregulated upon ZNF91&DNMT1-CRISPRi in NPCs (Figure 4E). One example was an SVA-E element located upstream of the RNF24 gene (Figure 4F). This SVA was transcriptionally silent in control iPSCs and control NPCs. In ZNF91-CRISPRi iPSCs we detected a robust activation of the expression of this SVA-E element that correlated with the loss of DNA methylation. When the ZNF91-CRISPRi iPSCs were differentiated to NPCs, the SVA remained expressed; this also correlated with a lack of DNA methylation. Thus, the loss of DNA methylation patterns over SVAs in iPSCs upon ZNF91-CRISPRi correlates with the transcriptional activation of SVAs, including when these cells are differentiated to NPCs. These experiments demonstrate that ZNF91 dynamically represses the expression of at least some SVAs in iPSCs, and is essential for establishing stable transcriptional repression of these SVAs.

### DNA methylation and H3K9me3 co-operate to protect the human genome from the cis-regulatory influence of SVAs

SVAs carry regulatory sequences which can mediate *cis*-acting transcriptional effects on the surrounding genome ^26–31^. We therefore investigated whether the ZNF91-mediated heterochromatin domains found over SVAs in NPCs influenced this activity. When investigating transcriptional changes of genes monitored via RNA-seq upon removal of H3K9me3 (ZNF91-CRISPRi), removal of DNA methylation (DNMT1-KO), or removal of both repressive marks (ZNF91&DNMT1-CRISPRi) we only found profound effects on nearby gene expression in the ZNF91&DNMT1-CRISPRi NPCs. The expression of genes located in the vicinity of an SVA element were significantly increased upon DNMT1&ZNF91-CRISPRi but not when deleting only one of the factors (Figure 5A). This effect could be detected when the SVA was located up to 50 kbp from the TSS, but was stronger when the SVA was closer to the TSS (Figure 5A).

**Figure 5.**
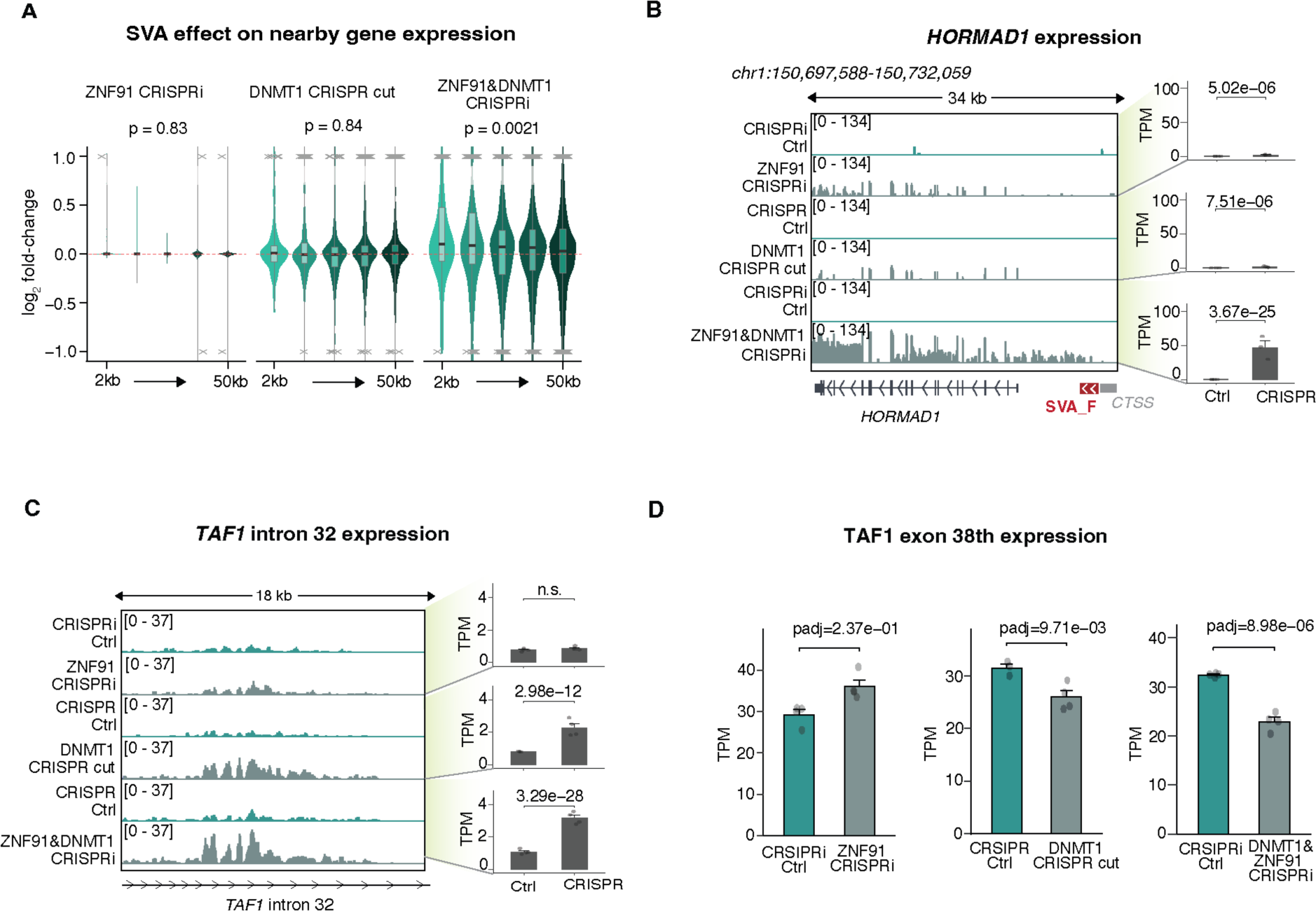
SVAs have a regulatory influence on nearby genes when heterochromatin marks are lost. (A) Violin plot showing the effect of SVAs on nearby gene expression (2-50 kbp) in ZNF91-CRISPRi, DNMT1-KO, and ZNF91&DNMT1-CRISPRi (One-way ANOVA). (B) Genome browser tracks (left) showing *HORMAD1* expression in ZNF91-CRISPRi, DNMT1-KO, and ZNF91&DNMT1-CRISPRi. Barplots (right) showing *HORMAD1* expression in ZNF91-CRISPRi, DNMT1-KO, and ZNF91&DNMT1-CRISPRi (padj, DESeq2). (C) Genome browser tracks (left) showing *TAF1* intron 32 expression in ZNF91-CRISPRi, DNMT1-KO and ZNF91&DNMT1-CRISPRi. Barplots (right) showing *TAF1* intron 32 expression in ZNF91-CRISPRi, DNMT1-KO and ZNF91&DNMT1-CRISPRi (padj, DESeq2). (D) Barplots showing *TAF1* exon 38 expression in ZNF91-CRISPRi, DNMT1 CRISPR-cut, and DNMT1&ZNF91-CRISPRi (padj, DESeq2).

Notably, the dynamics of the SVA-mediated influence on gene expression was distinct between different loci. Most genes in the vicinity of an SVA were completely unaffected by ZNF91 deletion or DNMT1 deletion alone, but transcriptionally upregulated when both factors were removed (Figure 5A). Thus, in most instances the presence of one of the heterochromatin marks was sufficient to protect flanking genomic regions from the regulatory impact of SVAs. However, we also found examples where both marks were needed to block the regulatory impact of SVAs. For example, the expression of *HORMAD1* was upregulated due to the activation of an upstream SVA-F element acting as an alternative promoter in both ZNF91-CRISPRi and DNMT1-KO NPCs (Figure 5B). When both ZNF91 and DNMT1 were inhibited, *HORMAD1* expression was even more strongly activated, suggesting a co-operative mode of action (Figure 5B). This demonstrates that at some loci both epigenetic marks are necessary to block the regulatory impact from SVAs at some loci.

### Loss of H3K9me3 and DNA methylation over the XDP-SVA results in an aggravated molecular phenotype at the TAF1 locus

We next used the XDP-NPCs to investigate if the presence of H3K9me3 and DNA methylation over the XDP-SVA has any impact on *TAF1* expression. Removing H3K9me3 alone (ZNF91-CRISPRi) did not affect intron retention in the *TAF1* loci in the XDP-NPCs nor exon 38 expression of the *TAF1* gene (Figure 5C, D). When we investigated the *TAF1* loci in XDP-NPCs that lacked DNA methylation (DNMT1-KO), retention of intron 32 was significantly increased and exon 38 expression was reduced (Figure 5C, D). Removing both DNA methylation and H3K9me3 (ZNF91&DNMT1-CRISPRi) had an even stronger effect on *TAF1* expression, including a considerable increase in intron 32 retention of *TAF1* and lower expression of exon 38 in XDP-NPCs (Figure 5C, D). Thus, the loss of both DNA methylation and H3K9me3 aggravates the molecular pathology in XDP-NPCs. These data demonstrate that the regulatory impact of the polymorphic XDP-SVA is negatively influenced by the presence of a local mini-heterochromatin domain. When this heterochromatin domain is lost, the *cis*-regulatory effect of the XDP-SVA is strongly and significantly enhanced.

### Polymorphic SVA insertions are silenced by ZNF91/H3K9me3 and DNA methylation

To investigate whether the local heterochromatin observed over the polymorphic XDP-SVA represented a unique event or if it was a general effect, we extended our analysis to other polymorphic SVAs in the genomes of two of the individuals in this study. We took advantage of the whole genome ONT long-read sequencing data from the XNPC1 and CNPC1 lines and used the Transposons from Long DNA Reads (TLDR) pipeline to identify non-reference SVA insertions (Figure 6A) ^61^. We identified 22 high-confidence polymorphic insertions of the SVA-E and -F subfamilies (average 2.5 kbp in length, range 1.23.8 kbp), of which 14 were shared between the two genomes (Figure 6B). Notably, several of these polymorphic SVAs, which represent recent TE insertions into the germline of these two individuals, displayed clear hallmarks of local heterochromatinization, including the presence of DNA methylation and H3K9me3.

**Figure 6.**
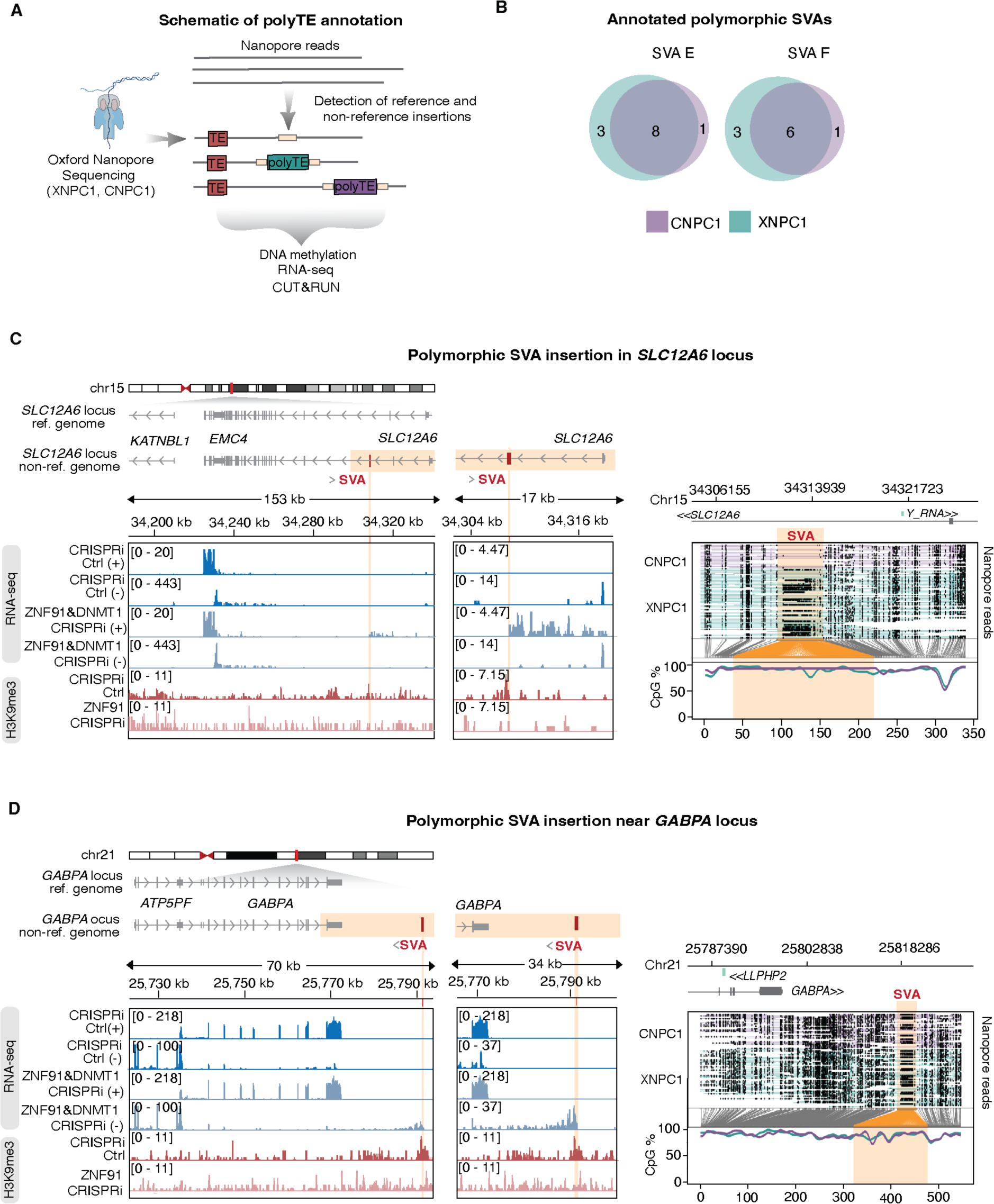
Polymorphic SVA insertions are repressed by DNA methylation and H3K9me3. (A) Schematic of ONT sequencing and annotation of polymorphic SVAs. (B) Venn diagram showing annotated polymorphic SVA insertions. (C) Genome browser tracks (left) showing gene expression and H3K9me3 over a polymorphic SVA insertion and the nearby gene *SLC12A6*. ONT sequencing data (right) showing DNA methylation over the annotated polymorphic insertion. (D) Genome browser tracks (left) showing gene expression and H3K9me3 over a polymorphic SVA insertion and the nearby gene *GABPA*. ONT sequencing data (right) showing DNA methylation over the annotated polymorphic insertion.

For example, we found a polymorphic SVA insertion present only in XNPC1 in an intron of *SLC12A6*, which is a gene encoding a potassium/chloride transporter linked to neurological disorders ^81,82^. This SVA insertion site displayed H3K9me3 at its boundaries and was fully covered by DNA methylation (Figure 6C). ZNF91-CRISPRi led to loss of H3K9me3 over this SVA, while ZNF91&DNMT1-CRISPRi in XNPC1 led to its transcriptional activation, resulting in the expression of an antisense readthrough transcript extending into the *SLC12A6* gene (Figure 6C). Another example was a polymorphic SVA insertion shared between CNPC1 and XNPC1 downstream of *GABPA*, which is a gene encoding a DNA binding protein involved in mitochondrial function ^83,84^. Similarly, the SVA insertion results in the accumulation of H3K9me3 and DNA methylation; ZNF91-CRISPRi and ZNF91&DNMT1-CRISPRi resulted in the loss of H3K9me3 and transcriptional activation of the SVA respectively, generating a readthrough antisense transcript (Figure 6D).

These data confirm that recent, polymorphic germline SVA insertions are recognized by ZNF91 in human NPCs and are covered by a dual layer of repressive epigenetic marks. Loss of this local heterochromatin domain results in transcriptional activation of these elements and the formation of novel transcripts, which are likely to have a regulatory impact on nearby genes. In these cases this is illustrated by the production of polymorphic antisense transcripts to *SLC12A6* and *GABPA*.

## Discussion

Our data support a model in which the KZNF ZNF91 binds to SVAs in early human development and throughout brain development, here modeled using iPSCs and NPCs. ZNF91 binding results in the establishment of local heterochromatin over SVAs, characterized by DNA methylation and H3K9me3, in a TRIM28-dependent manner. In early development (here represented by iPSCs) both modifications are dependent on the binding of ZNF91 to SVAs. In later phases of development, after epigenetic reprogramming of the genome (here represented by NPCs) only H3K9me3 depends on ZNF91: DNA methylation over SVAs is propagated by DNMT1. These two repressive chromatin modifications work together to limit the influence of SVAs on the host genome. They prevent the expression of the SVA elements and restrict the *cis*-acting influence of SVAs on the surrounding regions in neural cells. It is worth noting that this mechanism does not only involve evolutionarily older SVA insertions that are fixed in the human population, but also include recent polymorphic germline SVA insertions including the disease-causing XDP-SVA.

SVAs carry regulatory sequences with the potential to provide strong *cis*-acting influences on gene regulatory networks ^23,26–31^. The ZNF91-mediated mini-heterochromatin domains prevent this *cis* influence in the NPC model system. Our data demonstrate that for most SVA loci only one of the heterochromatin marks is necessary to silence SVA expression and to prevent its regulatory influence on nearby gene expression. However, there are examples where the cooperation of the two mechanisms appears necessary. The most striking example is the *HORMAD1* locus, where an upstream SVA can act as an alternative promoter ^53^. The regulatory effect of this SVA is activated when H3K9me3 and DNA methylation are removed individually, demonstrating the need for both marks to prevent the *cis* influence of this SVA. Removal of both marks results in a massive activation of *HORMAD1* expression, indicating that dual removal has synergistic effects. It is not yet understood why some SVA loci, such as the one upstream of *HORMAD1*, require both mechanisms for their control. However, it is likely that the transcriptional and epigenetic state of the integration sites is important, as well as structural variants within SVAs. For example, it is known that the VNTR region of SVAs is highly variable and has expanded recently in human evolution ^61,85^. It is also evident that ZNF91-heterochromatin domains are not able to prevent the regulatory influence of all new SVA germline insertions. SVA insertions on the sense strand within genes are less abundant than expected by chance, suggesting that such SVA insertions are selected against ^16,86^. In addition, there is a growing number of polymorphic SVA insertions linked to genetic disorders, with XDP being the best characterized example.

The SVA insertion linked to XDP is in intron 32 of the essential gene *TAF1*; the molecular phenotype includes intron retention and reduced TAF1 expression ^39,40^. Our data demonstrate that ZNF91 binds to the XDP-SVA and establish a polymorphic mini-heterochromatin domain. The epigenetic status of the XDP-SVA is highly relevant for XDP pathology, as this layer of heterochromatin protects against the gene regulatory impact of the SVA. When DNA methylation and H3K9me3 are lost, intron retention is greatly increased, and *TAF1* expression levels are further reduced. Thus, while ZNF91 can limit the impact of the XDP-SVA insertion, it cannot entirely remove the regulatory impact over the *TAF1* gene. This explains why the XDP-SVA causes disease while most other SVA insertions are inert. However, we still do not understand why ZNF91 is unable to fully block the *cis*-regulatory impact of the XDP-SVA.

Our data are limited to cell-culture models; the epigenetic status of the XDP-SVA in human brain tissue has not been investigated. It is worth noting that DNA methylation patterns in the human brain change with age ^87–89^. It will be interesting to investigate if DNA methylation over the XDP-SVA is stable in the human brain, or if it is lost with ageing. Such phenomena could explain the late-onset phenotype of XDP, where a gradual increase in the loss of *TAF1* function ultimately results in cellular dysfunction. Such a scenario would also open new therapeutic possibilities where restoration of DNA methylation on the XDP-SVA could block or reverse the pathology.

KZFPs have been implicated in an evolutionary arms race with TEs, where KZFP gene expansions and modifications limit the activity of newly-emerged transposon classes ^50,90^. This event is followed by mutations in the TEs to avoid repression in an ongoing cycle. One such example is ZNF91, which appeared in the last common ancestor of humans and Old-World monkeys and underwent a series of structural changes about 8-12 million years ago that enabled it to bind to SVA elements ^50,53^. However, the presence of the SVA-binding ZNF91 in hominoids has not prevented the expansion of SVAs in their genomes. On the contrary, SVAs are highly active in the human germline, providing a substantial source of genome variation in the population ^5,10,21–25^. Although ZNF91 is not able to completely prevent new SVA germline insertions, sustained expression during brain development greatly limits the *cis*-regulatory impact of these insertions. Thus, it appears that in this case ZNF91 may facilitate the expansion of SVA insertions by limiting their gene-regulatory impact on the human genome. Thus, our data are consistent with a model where KZFPs are not only TE repressors, but also facilitators of inert germline transposition events, thereby fueling genome complexity and evolution.

Our results demonstrate how a KZFP prevents the regulatory impact of TEs in human neural cells, but it is still not known if the ZNF91-SVA partnership represents a unique event, or if these results can be extrapolated to other TE families and lineages. In humans, there are three active TE classes: Alus, LINE-1s, and SVAs. The relationship between LINE-1s and SVAs is of special interest, since SVA retrotransposition depends on co-expression of the LINE-1 machinery. We and others have previously found that LINE-1s are controlled by DNA methylation in human NPCs, and that their transcriptional activation is recognized and silenced by the HUSH complex by a mechanism that is independent of KZFPs and TRIM28 ^59,91–93^. Thus, in human brain development LINE-1 activity appears to be controlled by fundamentally different mechanisms to SVAs. This suggests that the control of TE activity in the human brain is not only multilayered, but also highly specialized. Our data are limited to cell models of early development and neural cells, where *ZNF91* is particularly highly expressed. We do not know how SVAs are controlled in other human tissues. In addition, our data indicate that there are additional KZFPs controlling SVAs in early human development (see one example in Figure S4). ZNF91 deletion in iPSCs activates only a fraction of SVAs, all others stayed silenced. It is likely that these SVAs contain binding sites for additional KZFPs, such as ZNF611, that co-operate to control SVAs in early development ^29,53^. ZNF91 may then play a unique role in neural tissues. It is the only KZFP which binds SVAs and protects against *cis*-acting mechanisms from regulatory sequences in SVAs that are highly active in neural cells.

In summary, our results provide a unique mechanistic insight into an epigenetic defense system, based on a KZFP, active against the regulatory impact of SVA transposons in the human brain. On the one hand, this system protects the genome from any negative impact of SVAs, and SVA insertions result in genetic disease only in very rare instances, here exemplified by XDP. On the other hand, this system has likely contributed to the expansion of SVAs in our genomes, maximizing the potential for TEs to contribute to increased genome complexity and suggesting that SVAs are likely to have played an important role in primate brain evolution.

### Data and code availability

Processed sequencing data has been deposited at GSE245093. Additional information required to reanalyze the data reported in this paper is available from the lead contact upon request.

This paper includes analyses of existing, publicly available data. The accession numbers for these datasets are:

GSE224747: 3’ single nuclei RNAseq, bulk RNAseq, and H3K9me3 CUT&RUN of human fetal forebrain tissue. GSE242143: H3K9me3 CUT&RUN from the DNMT1-KO NPCs

All original code has been deposited at GitHub and is publicly available at: https://github.com/raquelgarza/XDP_Horvath_2023

## Acknowledgements

We would like to thank Frank Jacobs, Didier Trono, Cristopher Bragg, and Amy Alessi for comments on the manuscript and their support throughout this project. We also thank Jenny Johansson, Marie Persson Vejgården, Anna Hammarberg, and Ulla Jarl for their technical assistance. We are grateful to all members of the Jakobsson lab. The work was supported by grants from the Collaborative Center for X-Linked Dystonia-Parkinsonism (J.J and C.H.D.), the Swedish Research Council (2018-02694 to J.J. and 2021-03494 to C.H.D.), the Swedish Brain Foundation (FO2019-0098 to J.J.), Cancerfonden (190326 to J.J.), Barncancerfonden (PR2017-0053 to J.J.), the Swedish Society for Medical Research (S19-0100 to C.H.D.), and the Swedish Government Initiative for Strategic Research Areas (MultiPark & StemTherapy).

## Author Contributions

All of the authors took part in designing the study and interpreting the data. V.H. and J.J. conceived the study. V.H., M.J., P.J., A.A., G.C., O.K., L.C.V. and C.D. performed the experimental research. R.G. and N.P., performed the bioinformatic analyses. C.D., P.G. contributed expertise. V.H., C.D. and J.J. wrote the manuscript, and all authors reviewed the final version.

## METHODS

### EXPERIMENTAL MODEL AND SUBJECT DETAILS

Prior to experimental use, all cell lines were confirmed to be mycoplasma free.

#### Induced pluripotent stem cells (iPSCs)

We used iPSC lines derived from three XDP patients and three healthy individuals from WiCell (See table 1). iPSCs were maintained on Biolaminin 521-coated (0.7 mg/cm^2^; Biolamina) Nunc multidishes in iPS media (StemMACS iPS-Brew XF and 0.5% penicillin/streptomycin (GIBCO)). Cells were passaged 1:3 every 2-3 days. Briefly, cells were rinsed once with DPBS (GIBCO) and dissociated using Accutase (GIBCO) at 37°C for 5 minutes. Following incubation, Accutase was carefully aspirated from the well, and the cells were washed off from the dish using washing medium (9.5 ml DMEM/F12 (GIBCO) and 0.5 ml knockout serum replacement (GIBCO)). The cells were then centrifuged at 400 g for 5 minutes and resuspended in iPS brew medium supplemented with 10 mM Y27632 Rock inhibitor (Miltenyi Biotech) for expansion. The media was changed daily ^95^.

#### Neural progenitor cells (NPCs)

The neural progenitor cells (NPCs) were generated from iPSCs from the three XDP patients and three unaffected individuals (Table 1). The neural induction was done as described in ^96^. The NPCs were cultured in DMEM/F12 (Thermo Fisher Scientific) supplemented with glutamine (2 mM, Sigma), penicillin/streptomycin (1×, Gibco), N2 supplement (1×, Thermo Fisher Scientific), B27 (0.05×, Thermo Fisher Scientific), EGF and FGF2 (both 10 ng/ml, Thermo Fisher Scientific). 10 mM Y27632 Rock inhibitor (Miltenyi) was also used. Cells were grown on Nunc multidishes or in T25 flasks pre-coated with Poly L-Ornithine (15 μg/ml, Sigma) and Laminin (2 μg/ml, Sigma). Cells were passaged every 2-3 days using TryplLE^™^ express enzyme (GIBCO) and trypsin inhibitor (GIBCO).

#### Immunocytochemistry

24-well Nunc plates were pre-coated with Poly L-Ornithine (15 μg/ml, Sigma) and Laminin (2 μg/ml, Sigma). Approximately 50,000 cells were plated in the wells and were allowed to expand, until they reached 70-80% confluency. At this point, cells were washed three times with DPBS (GIBCO) and fixed with 4% paraformaldehyde (Merck Millipore) solution for 15 minutes at room temperature, and washed again three times with DPBS. Fixed cells were stored in DPBS at 4°C for a maximum of one month until staining and imaging.

For blocking, cells were incubated for one hour with 5% normal donkey serum (NDS) in TKBPS (KBPS with 0.25% Triton X-100 (Fisher Scientific)). Subsequently, they were incubated overnight at 4°C with the primary antibody (5mC, Active Motif, cat.no. 39649, lot 02617020, used 1:250; SOX2, R&D Systems, AF2018, 1:100; and Nestin, Abcam, AB176571, 1:100). For a negative control, cells were incubated overnight with TKPBS + 5% NDS. After overnight incubation, cells were washed two times for five minutes in TKPS, followed by five minutes in TKBPS with NDS. Next, they were incubated at room temperature for two hours with the secondary antibody (donkey anti-rabbit Alexa fluor 647, 1:200, Jackson Lab, and donkey anti-goat cy3, 1:200, Jackson Lab) and for five minutes with DAPI (1:1000, Sigma Aldrich) as a nuclear counterstain. This was followed by two five-minute washes with KPBS, then cells were stored in PBS until imaging.

##### 5mC staining

As described in Jönsson et al. 2019 ^59^, cells stained for 5mC were pre-treated with 0.9% Triton in PBS for 15 minutes, followed by 2 N HCl for 15 minutes, then 10 mM Tris-HCl, pH 8, for 10 minutes prior to incubation with the primary antibody. Cells were imaged using a fluorescence microscope (Leica).

#### RNA sequencing

Total RNA was isolated using the RNeasy Mini Kit (Quiagen) with on-column DNAse treatment following the manufacturer’s instructions. The isolated RNA was used for qPCRs (see below) and RNA sequencing. RNA sequencing was performed using four biological replicates. Libraries for RNA sequencing were generated using Illumina TruSeq Stranded mRNA library prep kit (poly-A selection), optimized for long fragments, and sequenced on a Novaseq6000 (paired end, ∼250 bp) yielding an average of 46M reads. The reads were mapped to the human reference genome (hg38) using STAR aligner v2.7.8a ^97^ and gene quantification was performed using FeatureCounts (Subread package v1.6.3; hg38 Gencode v38), setting -p for paired-end, and -s 2 for reversely stranded reads (TruSeq) ^98^.

To quantify TE expression, reads were re-mapped using STAR aligner and discarded if mapped to more than one location (–outFilterMultimapNmax 1). A maximum of 0.03 mismatches per base were allowed (–outFilterMismatchNoverLmax 0.03). FeatureCounts (Subread package v1.6.3) using hg38 RepeatMasker annotation “parsed to filter out low complexity and simple repeats, rRNA, scRNA, snRNA, srpRNA and tRNA” was used to quantify reads ^99^.

Bigwig files for genome browser tracks were generated using bamCoverage (deeptools v2.5.4), set to –normalizeUsingRPKM and –filterRNAstrand to split signal between strands. Visualization was performed in the Integrative Genome Browser (IGV) ^100^. Matrices for deeptools heatmaps were generated including only SVAs longer than 1 kbp (grouped by subfamily; individual BED files), using computeMatrix scale-regions setting –regionBodyLength to 1kbp, and flanking regions (-a and -b) to 10kbp. Heatmaps were generated using plotHeatmap (v3.5.1). Profile plots were generated the same way, ungrouping the SVAs (input a single BED file) prior to the matrix computation.

Normalization of counts to visualize the expression of different features on barplots (genes, *TAF1* intron 32 or exon 38) was performed as TPM: the length of the feature was used to calculate an approximate TPM value. Statistical tests, however, were performed using DESeq2, which normalizes using median of ratios ^101^. Intron 32 and exon 38 of the *TAF1* gene were added as part of the gene count matrix.

### CUT&RUN

CUT&RUN analysis was done on CNPC1, CNPC2, XNPC1, and XNPC3 in both ZNF91-CRISPRi and TRIM28-CRISPRi, including CRISPRi-Ctrl (lacZ). We followed the protocol described in Skene and Henikoff 2018 ^102^. Briefly, 300,000 cells were washed twice (20 mM HEPES pH 7.5, 150 mM NaCl, 0.5 mM spermidine, 1× Roche cOmplete protease inhibitors) and attached to 10 ConA-coated magnetic beads (Bangs Laboratories) that had been pre-activated in binding buffer (20 mM HEPES pH 7.9, 10 mM KCl, 1 mM CaCl_2_, 1 mM MnCl_2_). Bead-bound cells were resuspended in 50 ml buffer (20 mM HEPES pH 7.5, 0.15 M NaCl, 0.5 mM spermidine, 1× Roche complete protease inhibitors, 0.02% w/v digitonin, 2 mM EDTA) containing primary antibody (rabbit anti H3K9me3, Abcam ab8898, RRID:AB_306848, or goat anti-rabbit IgG, Abcam ab97047, RRID:AB_10681025) at 1:50 dilution and incubated at 4°C overnight with gentle shaking. Beads were washed thoroughly with digitonin buffer (20 mM HEPES pH 7.5, 150 mM NaCl, 0.5 mM spermidine, 1× Roche complete protease inhibitors, 0.02% digitonin). After the final wash, pA-MNase (a generous gift from Steve Henikoff) was added in digitonin buffer and incubated with the cells at 4°C for 1 hour. Bead-bound cells were washed twice, resuspended in 100 ml digitonin buffer, and chilled to 0-2°C. Genome cleavage was stimulated by adding 2 mM CaCl_2_ at 0°C for 30 minutes. The reaction was quenched by adding 100 ml 2× stop buffer (0.35 M NaCl, 20 mM EDTA, 4 mM EGTA, 0.02% digitonin, 50 ng/ml glycogen, 50 ng/ml Rnase A, 10 fg/ml yeast spike-in DNA (a generous gift from Steve Henikoff) and vortexing. After 10 minutes incubation at 37°C to release genomic fragments, cells and beads were pelleted by centrifugation (16,000 g, 5 minutes, 4°C) and fragments from the supernatant were purified. Illumina sequencing libraries were prepared using the Hyperprep kit (KAPA) with unique dual-indexed adapters (KAPA), pooled, and sequenced on a Nextseq500 instrument (Illumina). Paired-end reads (2×75) were aligned to the human and yeast genomes (hg38 and R64-1-1 respectively) using bowtie2 (–local –very-sensitive-local –no-mixed –no-discordant –phred33 -I 10 -X 700) and converted to bam files using samtools ^103^. Normalized bigwig coverage tracks were made using bamCoverage (deeptools) ^104^, with a scaling factor accounting for the number of reads arising from the spike-in yeast DNA (10^4^ per aligned yeast read number). Tracks were displayed in IGV.

#### CRISPR approaches

##### CRISPRi

To silence the transcription of *ZNF91* and *TRIM28*, we used the catalytically inactive Cas9 (deadCas9) fused to the transcriptional repressor KRAB ^105^. Single-guide sequences were designed to recognize DNA regions just down-stream of the transcription start site (TSS), according to the GPP Portal (Broad Institute). ZNF91 sgRNA: GAGTTTCCAGGTCTCGACTT (No PAM). The guides were inserted into a deadCas9-KRAB-T2A-GFP lentiviral backbone containing both the guide RNA under the U6 promoter and dead-Cas9-KRAB and GFP under the Ubiquitin C promoter (pLV hU6-sgRNA hUbC-dCas9-KRAB-T2a-GFP, a gift from Charles Gersbach, Addgene plasmid #71237 RRID:Addgene_71237). The guides were inserted into the backbone using annealed oligos and the BsmBI cloning site. Lentiviruses were produced as described below, yielding titers of 10^8^-10^9^ TU/ml, which was determined using qRT– PCR. Control virus with a gRNA sequence absent from the human genome (LacZ) was also produced and used in all experiments. All lentiviral vectors were used with an MOI of 2.5 unless stated differently. GFP cells were FACS isolated (FACSAria, BD sciences) on day 10 at 10°C (reanalysis showed >97% purity) and pelleted at 400 g for 5 minutes, snap frozen on dry ice and stored at −80°C until RNA isolation. All groups were performed in 4 biological replicates unless indicated differently. Knock-down efficiency was validated using RNA sequencing.

##### DNMT1 CRISPR cut

LV.gRNA.CAS9-GFP vectors were used to target *DNMT1* ^79^ or LacZ (control) as described in ^59^. Lentiviral vectors were produced as described previously and had a titer of 10^8^-10^9^ TU/ml which was determined using qRT-PCR. hNPCs were transduced with an MOI of 10-15, allowed to expand for 10 days, and were FACS-sorted as described previously.

##### DNMT1 and ZNF91 double CRISPRi

We used a double-transduction method to do a double CRISPRi of *DNMT1* and *ZNF91*. We transduced *ZNF91* with the previously-mentioned deadCas9-KRAB-T2A-GFP lentivirus containing the dead Cas9 protein and a GFP. At the same time, the cells were transduced with lentivirus containing the pLV.U6BsmBI.EFS-NS.H2b-RFPW lentiviral backbone wihth the gRNA for *DNMT1* and mCherry as a marker but without deadCas9 to knock down *DNMT1*. DNMT1 sgRNA: TGCTGAAGCCTCCGAGATGC (no PAM). Double-positive (mCherry and GFP) cells were FACS sorted as previously described and stored at −80°C until RNA extraction.

#### Lentiviral vector production

Lentiviral vectors were produced according to Zufferey et al. ^106^. Briefly, HEK293T cells were grown to a confluency of 70-90% at the day of transfection for lentiviral production. We used third-generation packaging and envelop vectors (pMDL, psRev, and pMD2G), together with polyethyleneimine (PEI Polysciences PN 23966, in DPBS (Gibco)). The lentivirus was harvested 2 days after transfection. The supernatant was then collected, filtered, and centrifuged at 25,000 g for 1.5 hours at 4°C. The supernatant was removed from the tubes and the virus was resuspended in PBS and left at 4°C. The resulting lentivirus was aliquoted and stored at −80°C.

### CUT&RUN qRT-PCR

To identify whether the XDP-SVA was surrounded by an H3K9me3 mark, we designed a qPCR approach. Briefly, two primers were designed on the 5’ flanking region of the XDP-SVA: one in the flanking region and one in the SVA. As the positive control, primers were designed for a genomic region (hg38, chr5:141253464-141255143) known to be covered by H3K9me3. The CUT&RUN library was used as a template for amplification. The qRT-PCR was done with SYBR Green I Master (Roche) on a LightCycler 480 (Roche). The primer pairs used were: XDP-SVA forward (5’-3’): GAATGGTATATGTTTAGTTTTACA; XDP-SVA reverse (5’-3’): CATGACCCTGCCAAATCCCCCT; positive-control forward (5’-3’): AAATGGGAATTAAAATCAGTGAGGC; positive-control reverse (5’-3’): TTGACATATCATTAAGGGGGCA.

#### Oxford Nanopore sequencing

##### Whole-genome Nanopore sequencing

DNA was extracted from frozen pellets using the Nanobind HMW DNA Extraction kit (PacBio) following the manufacturer’s instructions. Final product was eluted in 100 μl of elution buffer provided in the kit. DNA concentration and quality were measured using Nanodrop and Qubit from the top, middle, and bottom of each tube. Only DNA with a quality of 260/280 1.8-2.0 and 260/230 2.0-2.2 was further processed. Whole-genome sequencing on samples XNPC1 and CNPC1 was done at the SciLife lab in Uppsala using SQK-LSK109 Ligation Sequencing kit (Oxford Nanopore Technologies) and FLO-PRO002 PromethION Flow Cell R9 Version, and was sequenced on a PromethION (Oxford Nanopore Technologies).

##### Cas9-targeted Nanopore sequencing

To target the XDP locus we used the Cas9 sequencing (SQK-CS9109) kit following the manufacturer’s instructions (Oxford Nanopore Technologies). To enrich for the fragment of interest, four previously-described guide RNAs were used ^107^. Briefly, two guides were designed upstream and two were designed downstream of the XDP-SVA insertion. The excision using these guides resulted in a 5.5 kbp product, including the XDP-SVA (2.6 kbp). 5 μg of DNA was used. Samples were sequenced on a MinION Mk1Mc using flow cell R9.4.1 (Oxford Nanopore Technologies). One flow cell per sample was used. For cas9-targeted enrichment we obtained 20770 reads from 3 samples (XNPC3 CRISPRi-Ctrl 7805 reads; XNPC3 DNMT1-KO 7581 reads; XNPC3 ZNF91-CRISPRi 4384 reads; samtools view -c BAM). The proportion of reads that mapped to the target was on average 1.25% (CRISPRi-Ctrl 52 reads; DNMT1-KO 171 reads; ZNF91-CRISPRi 36 reads; primary alignments over *TAF1* intron 32 only: samtools view -c -F 260 BAM chrX:71424238-71457170). XNPC3 CRISPRi Ctrl, ZNF91-CRISPRi, and DNMT1-KO samples were sequenced using the targeted approach.

Fastq files were indexed using the nanopolish index (v0.13.3) on default parameters^108^. To build an index of the XDP genome (with the XDP-SVA insertion), a consensus of the SVA sequences (two enrichment methods via PCR and CRISPR) as reported by Reyes et al ^107^ (https://github.com/nanopol/xdp_sva/blob/main/) was created using EMBOSS cons (v6.6.0.0) (http://emboss.open-bio.org/) resulting in a 2,638 bp long sequence.

A *TAF1* fasta file was generated using grep -w *TAF1* from hg38 gencode v38, and bedtools getfasta (v2.30.0) ^109^. The SVA consensus sequence and the *TAF1* sequence were then aligned using clustalw2 (v2.1). We observed that the breaking point between the sequence extracted by Reyes et al. 2022 and the reference genome’s sequence of *TAF1* occurred at nucleotide chrX:71,440,502 ^107^.

Fasta file of chrX was then chopped using bedtools getfasta using breaking points:

chrX 1 71440502

chrX 71440503 156040895

The three sequences (chrX:1-71440502, the SVA sequence, and chrX:71440503-156040895) were then stitched (concatenated) together. A new genome fasta file was created concatenating all hg38 chromosomes fasta files (except for chrX) and the XDP-chrX fasta file. A minimap2 (v2.24) index was then created using the Nanopore present (-x map-ont) ^110^. Mapping of the reads was performed using minimap2 (v2.24) using the Nanopore preset (-a -x map-ont) with the XDP genome index. BAM files were sorted and indexed using samtools (v1.16.1).

Polymorphic insertions were identified using TLDR (v1.2.2), using GRCh38.p13 as reference genome (-r) and a TE library (-e) including the consensus sequences for TE subfamilies: L1Ta, L1preTa, L1PA2, SVA A-F, and HERVK (sequences provided by TLDR developers) ^61^. Insertions were considered for further analysis if they were found in the two individuals analyzed (CNPC1 and XNPC1). Insertions were required to have an UnmapCover of at least 80% (percentage of insertion with TEsequence), have a sequence similarity (TEMatch) of at least 80% to the TE consensus sequence, and a minimum of three reads supporting it (SpanReads).

The local consensus sequences of the polymorphic insertions (as output from TLDR) were introduced to the reference genome. A custom script (add_polymorphic_insertions_fa.py) was used to read the TLDR output table, sort the polymorphic insertions from the end to the start of each chromosome, and perform the following operations for each of the chromosomes:

1. Read its fasta file (chr.fa)
2. Extract its sequence before and after the insertion using bedtools getfasta (-fi chr.fa -bed coordinates.bed), where coordinates.bed included two coordinates: One spanning from the beginning of the chromosome to the start of the insertion (as reported by TLDR), and the second spanning from the end of the insertion (as reported by TLDR) to the end of the chromosome.
3. Concatenate the sequences:

a. The chromosome before the insertion
b. The sequence of the polymorphic insertion
c. The chromosome after the insertion

And re-write the chromosome’s fasta with it.

This process was repeated for each polymorphic insertion, introducing them in order from end to start of each of the chromosomes. Similarly, an updated gene annotation GTF file (to fit the coordinates including all polymorphic insertions) was created using a custom script following the same logic (add_polymorphic_insertions.py).

The reads were re-mapped to the custom genome using an indexed version of the output fasta (output from add_polymorphic_insertions_fa.py) using minimap2 (index using -x map-ont; mapping using -a -x map-ont).

Methylation for each of the regions of interest was called using nanopolish call-methylation (v0.13.3) with the raw reads (-r), the alignment files to the custom genome (-b), and the custom genome’s fasta file as a reference (-g). Databases for methylartist were produced using methylartist db-nanopolish (v1.2.2), using the methylation calls files as input (default parameters). Specific loci were visualized using methylartist locus (v1.2.2) ^60^.

## SUPPLEMENTARY FIGURES

**Figure S1.**
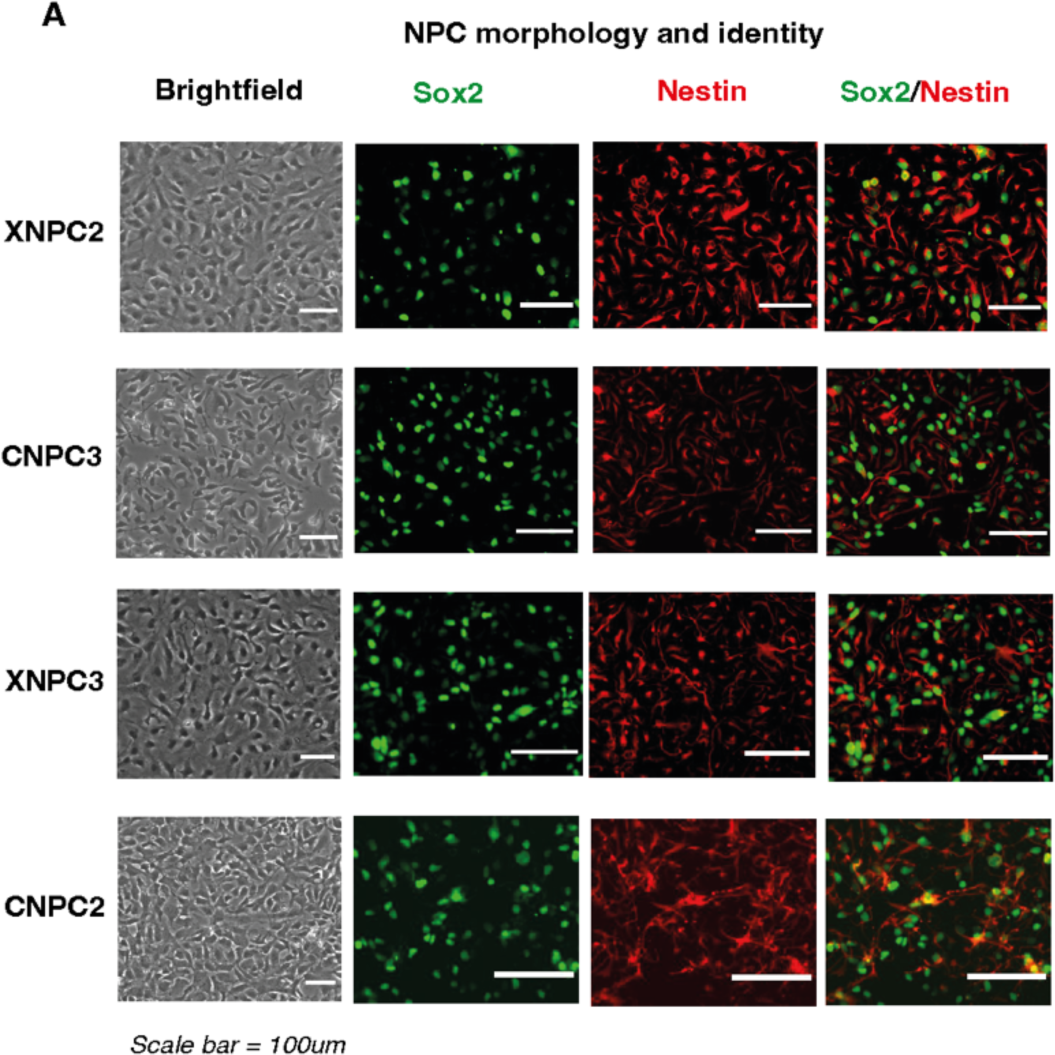
The XDP-NPCs display NPC morphology and expression of NPC markers. (A) Brightfield images of XDP-NPCs and Ctrl-NPCs (left). Immunostainings (right) of Sox2 (green) and Nestin (red) in XDP-NPCs and Ctrl-NPCs.

**Figure S2.**
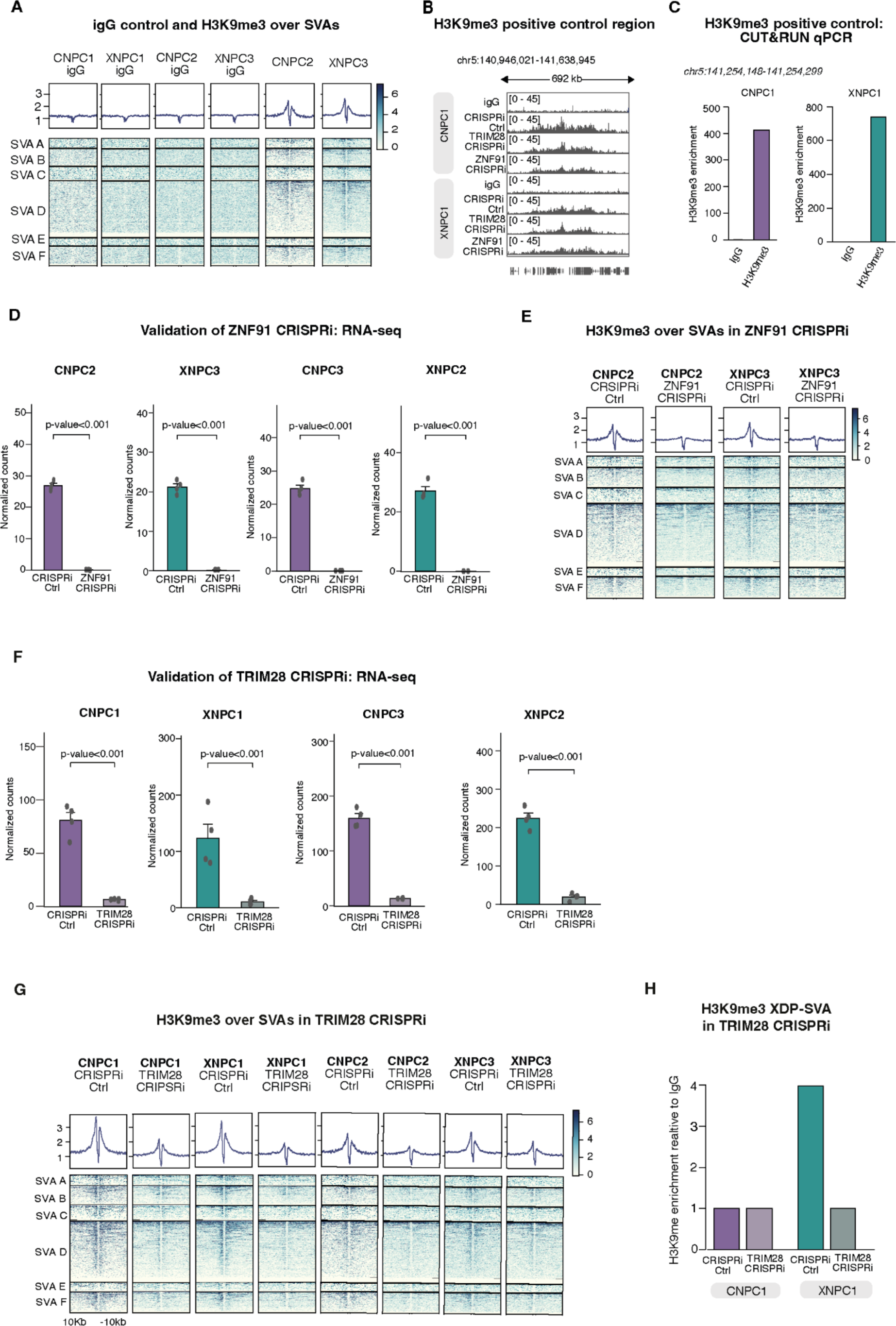
ZNF91 and TRIM28 orchestrate H3K9me3 deposition over SVAs in NPCs. (A) Heatmap showing igG and H3K9me3 enrichment in Ctrl-NPCs and XDP-NPCs. The genomic regions spanning ±10 kbp from the peak center are displayed. (B) Genome browser tracks showing H3K9me3 signal over a region known to be covered by H3K9me3 as a positive control. Tracks are shown for Ctrl-NPC and XDP-NPC for H3K9me3 and igG in control, TRIM28, and ZNF91-CRISPRi. (C) Barplots showing igG and H3K9me3 coverage over a positive-control region using CUT&RUN qPCR in Ctrl-NPC and XDP-NPC. (D) Barplots showing *ZNF91* expression (RNA-seq) in CRISPRi-Ctrl and ZNF91-CRISPRi (padj, DESeq2). (E) Heatmaps showing H3K9me3 signal around SVAs in CRISPRi-Ctrl and ZNF91-CRISPRi (n=2). (F) Barplots showing TRIM28 expression (RNA-seq) in CRISPRi-Ctrl and TRIM28-CRISPRi in Ctrl-NPC and XDP-NPC (n=4) (padj, DESeq2). (G) Heatmaps showing H3K9me3 signal around SVAs in CRISPRi-Ctrl and TRIM28-CRISPRi (n=4). (H) Barplots showing the H3K9me3 status of the XDP-SVA in CRISPRi-Ctrl and TRIM28-CRISPRi (n=2) (padj, DESeq2).

**Figure S3.**
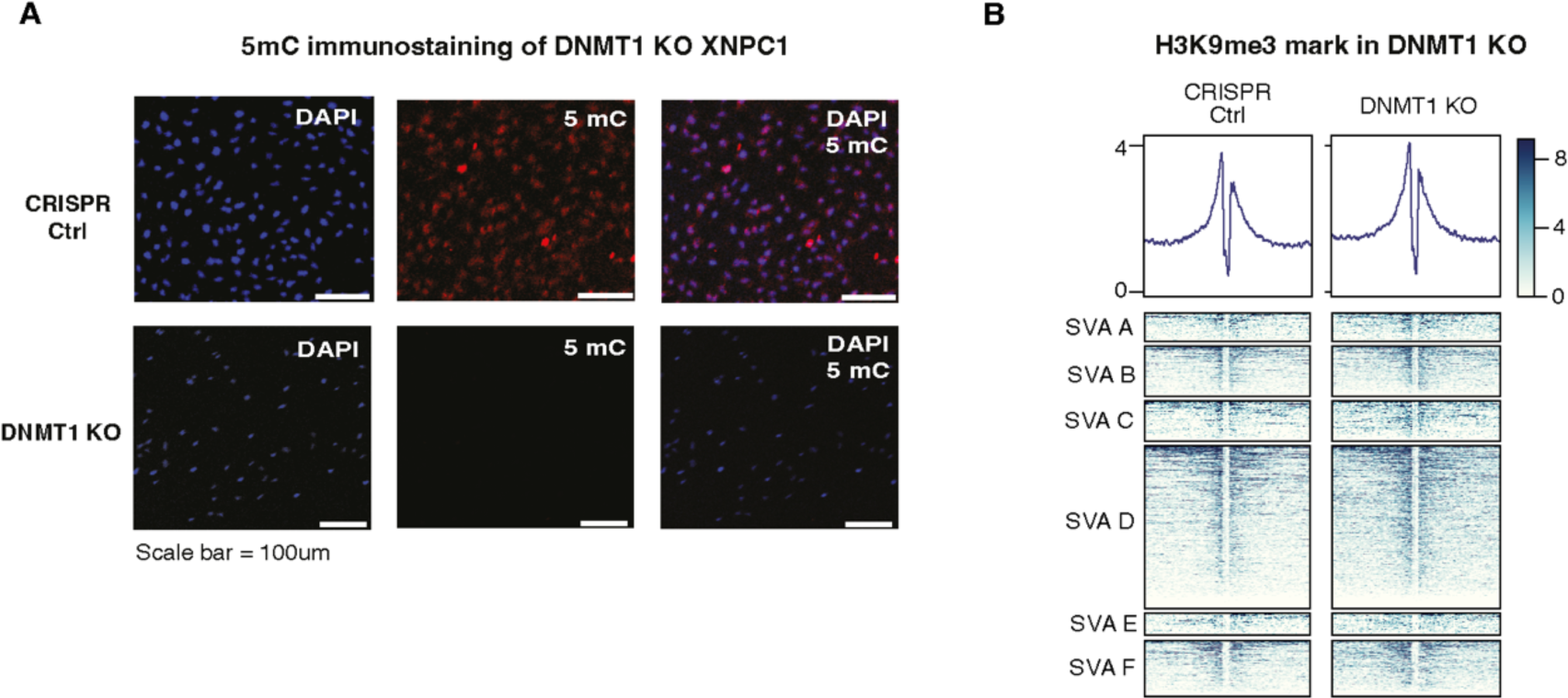
DNMT1 does not regulate H3K9me3. (A) Fluorescent 5mC immunostaining shows successful DNMT1-KO in XNPC1 10 days post transduction. Blue=Dapi, red=5mC. (B) Heatmap showing H3K9me3 around SVAs in a genome-wide scale in Ctrl and DNMT1-KO NPCs.

**Figure S4.**
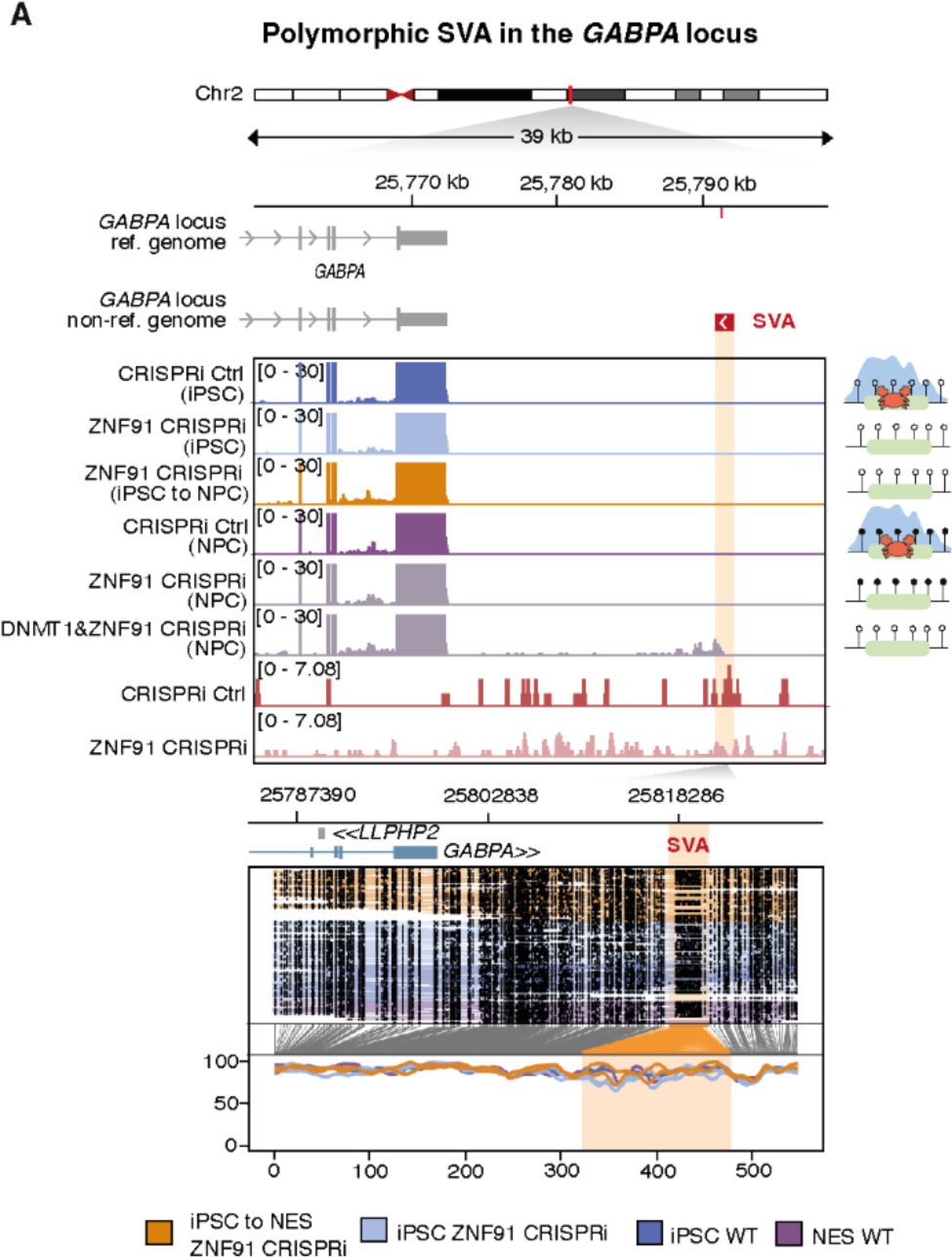
ZNF91 independent SVA regulation. (A) Polymorphic SVA loci near *GABPA.* Genome browser tracks (top) showing gene expression and H3K9me3. ONT reads (bottom) showing SVA DNA methylation.

